# Effect of predicting familiar melodies on alpha power

**DOI:** 10.1101/2023.03.16.532954

**Authors:** Shuma Ito, Kazuki Matsunaga, Ingon Chanpornpakdi, Toshihisa Tanaka

## Abstract

The processing of music and language share similar characteristics. Previous studies indicated that the similarity between language and melody was observed in event-related potentials to the deviation of a word and tone, respectively. We focused on a language study that demonstrated strong suppression of the alpha power in the presence of easily predictable words. Motivated by the physiological similarity between language and music, this study hypothesized that predictable music might suppress the alpha power. We measured electroencephalogram (EEG) signals while a melody followed by silence was presented for a participant who imagined the melody during the silent part of the music. The participants scored the melody’s familiarity to quantify the ease of prediction and imagination. We observed similarity to language processing. For familiar melodies, alpha power suppression was observed in the left frontal and left central regions. Further, we observed Bereitschaftspotential (a negative slope) in both familiar and unfamiliar conditions before the silent interval. Moreover, a network analysis revealed that information flow from the right sensory-motor cortex to the right auditory cortex in the beta band was stronger for familiar music than for unfamiliar music. Considering the previous findings that motor preparation and execution suppress the alpha power in the left frontal and left central regions, the alpha band suppression under music prediction suggests a motor interaction during music processing in the prediction of melodies.

## 1 Introduction

Music includes melody, rhythm, pitch, duration, and accent elements. Musical syntax is created using these elements, and music has a “context” like a language. The similarities between music and language have been studied in neuroscience.

The processing of language and melody is biased toward either the left or right hemisphere (lateralization) (Friederici and Alter, 2004; Kimura, 1964), and memory for language and melody occurs in the temporal lobes of both hemispheres (Booth et al., 2002; Satoh et al., 2006). Also, music processing and language display common features in the brain.

When we focus on the context of the melody, we can observe event-related potential (ERP) similar to that of language. Kutas and Federmeier (2000) reported that N400, a negative potential with a peak at approximately 400 ms after stimulus presentation, appeared when out of context language expression was presented. Miranda and Ullman (2007) measured the electroencephalogram (EEG) when presented with auditory stimuli that altered a note of a melody conditioned by familiarity and observed N400 under the condition that the given melody was familiar. N400 is related to memory access, such as semantic memory and recognition memory (Kutas and Federmeier, 2011). Therefore, memory access exhibits similar ERPs for language and melody stimuli.

In addition, alpha power is associated with the processing of music and language (Schaefer et al., 2011; Wang et al., 2018, 2022). In particular, alpha power suppression occurs for prediction from context (Rommers et al., 2017; Wang et al., 2018; Terporten et al., 2019; Gastaldon et al., 2020). Wang et al. (2018) requested participants to predict the next word in a context-sensitive task and found that magnetoencephalography (MEG) alpha power (8–12 Hz) was suppressed in the left frontal and left temporal lobes when the prediction was easier. Terporten et al. (2019) reported that predictability might be not only related to the strength of suppression in the alpha and beta bands but is also related to ERPs.

In the present study, we hypothesized that physiological mechanisms for prediction possess similarities and the brain processes music and language have many similarities. When conditioned on the predictability of a melody, we hypothesized that 1) ERPs occur when predicting melodies, regardless of the predictability, and 2) when the melody is more predictable, alpha power is more suppressed than when the melody is less predictable. We used music familiarity as a measure of the predictability of the melody. The reason for using familiarity is that it is more predictable when the music is familiar and less predictable when it is unfamiliar (Miranda and Ullman, 2007). To investigate these two hypotheses, we measured the EEG signals while performing a listening task; participants listening to a melody containing a silent section and predicting the missing notes. We confirmed the appearance of ERP by averaging the EEG data in the same way as Wang et al. (2018). We also confirmed the temporal fluctuation of alpha power by using time-frequency analysis. Furthermore, we quantified the connectivity between different regions by Phase Transfer Entropy (PTE). We also confirmed the frequency response of the EEG during the melody presentation and silent intervals using frequency analysis.

## 2 Materials and Methods

### 2.1 Participants

Twenty healthy young adults (men: 10 women: 10; age: 22±1.5 [range: 20–27] years) participated in this study. All participants self-reported that they possessed normal hearing and had no musical training. Some participants were lab students. Non-lab participants received compensetion of JPY 1,000 per hour. None of the student-participants were encouraged to participate by their professors, nor did they obtain any credits for doing so. All participants provided written informed consent. This study was approved by the Research Ethics Committee of Tokyo University of Agriculture and Technology.

### 2.2 Stimuli

EEG was measured during melody prediction using auditory stimulus comprising music with a beep at 440 Hz and a silent interval. We created 120 trials, three for each song, using 20 familiar and 20 unfamiliar songs chosen from the music used by Kumagai et al. (2018) and Miranda and Ullman (2007). We created music stimuli using Sibelius (Avid Technology, USA) and Python. We created all audio files with a sampling frequency of 44,100 Hz. We set the tempo of the melody at 2.5 Hz (Kumagai et al., 2018). A beep (duration 100 ms, pitch 440 Hz) was included before the melody to synchronize the presented melody with the EEG.

We placed a silent section in the middle of the melody and requested the participants to predict the melody during the silence section. Figure 1 shows an example of the melody. The silent section was set for 3 s between 8 and 11 s after the start of the melody. Each music stimulus duration of 20 s, including the beep.

**Figure 1:**
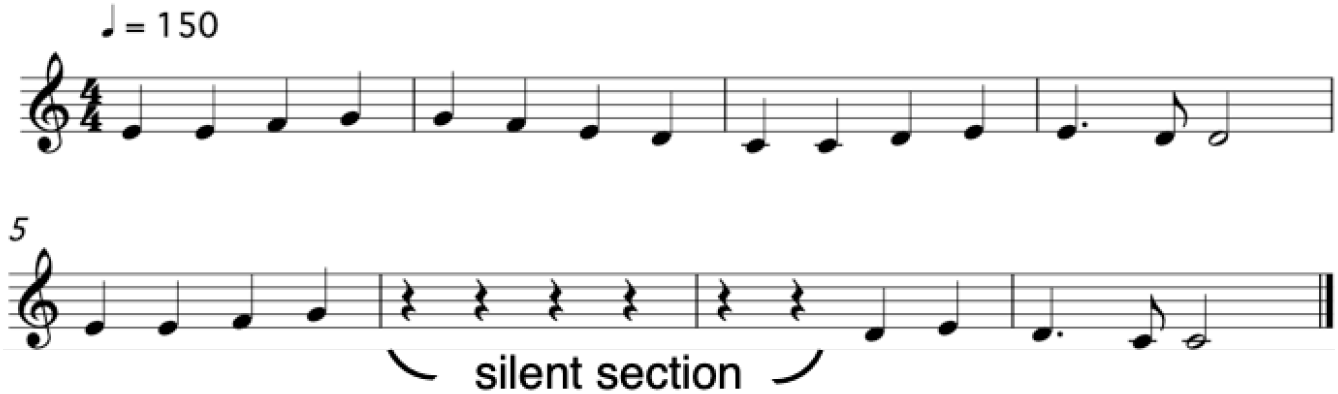
Example of the stimuli used in the experiment

### 2.3 Task

The participants performed the listening task by listening to a melody containing a silent section and predicting the missing notes. At the end of each melody, the participants answered whether they were familiar with the presented melody. We asked the participants to answer 5-level familiarity shown on the display with the mouse.

The task was performed in a soundproof room to prevent the influence of outside noise. In addition, we instructed the participants to fix their heads on the chin rest and retain the same position with minimal movement throughout the experiment. Furthermore, we instructed the participants to keep looking at the viewpoint shown on display during the presentation of the music stimuli.

The stimuli were presented using a loudspeaker (NS-F210, YAMAHA, Japan). The loudspeakers were placed at 92 ± 30°,and 1.3 m from the participants (Bregman, 1994). We used a monitor (VX279, ASUS, Taiwan) to display the instructions.

### 2.4 Procedure

Before initiating the task, the participants were allowed to practice. The participants listened to a melody with a silent section and practiced evaluating their familiarity with it. At this time, we confirmed that the participants understood the task correctly.

After completing the task practice, we initiated the experimental task as shown in Figure 2. A beep sounded 1.6 s after the task’s start, and the melody was presented after 1 s (Kong et al., 2014). The melody was presented and subsequently the participants answered whether they were familiar with it. Completion of the flow illustrated in Figure 2 was defined as one trial. There was a 3-s silent section from 8 to 11 s after the start of the melody. A total of 120 trials were conducted in 10 blocks of 12 trials each, with a 2-min rest between the blocks.

**Figure 2:**
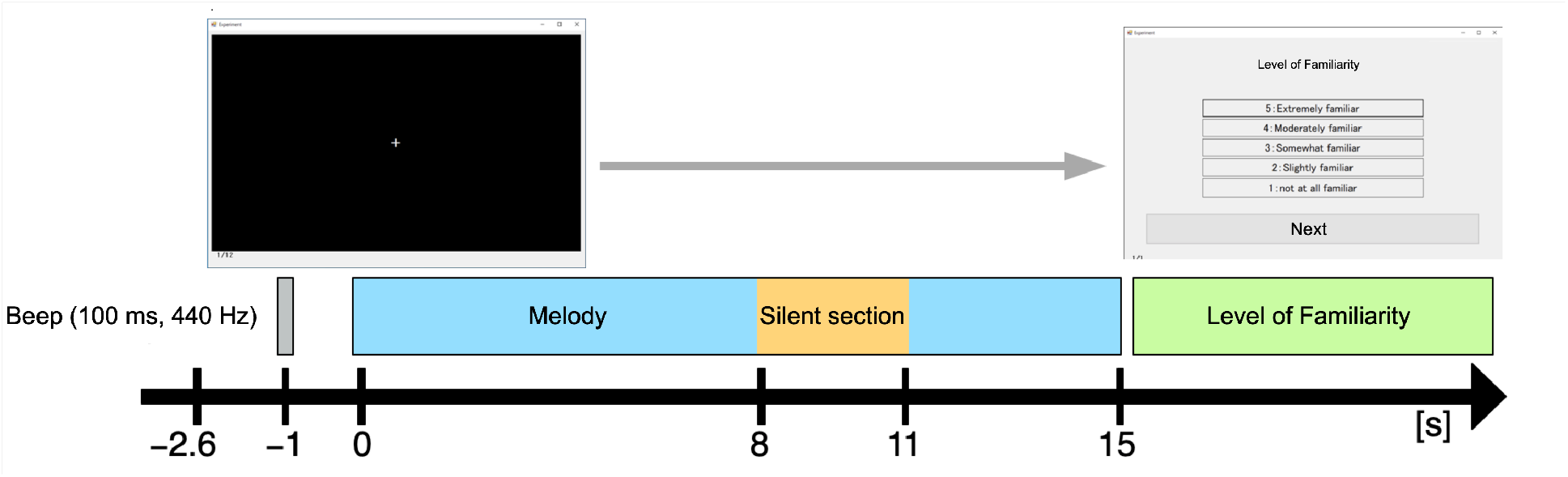
Experimental flow.

We evaluated familiarity with the melody using a Likert scale (Vagias, 2006). The rating was on a 5-point scale (5: Extremely familiar, 4: Moderately familiar, 3: Somewhat familiar, 2: Slightly familiar, 1: Not at all familiar).

### 2.5 Data Recording

EEG signals were measured using 64 scalp Ag/AgCl passive electrodes mounted in an EEG gel head cap (TMSi; Twente Medical Systems International, Oldenzaal, the Netherlands) and based on the international 10–10 system (Fp1, Fpz, Fp2, F7, F5, F1, F3, Fz, F2, F4, F6, F8, FC5, FC3, FC1, FCz, FC2, FC4, FC6, M1, T3, C5, C3, C1, Cz, C2, C4, C6, T4, M2, CP5, CP3, CP1, CPz, CP2, CP4, CP6, T5, T6, P5, P3, P1, Pz, P2, P4, P6, O1, Oz, O2, AF7, AF3, AF4, AF8, PO7, PO5, PO3, POz, PO4, PO6, PO8, FT7, FT8, TP7, and TP8), as shown in Figure 3. The electrodes’ impedance was kept below 10 kΩ.

**Figure 3:**
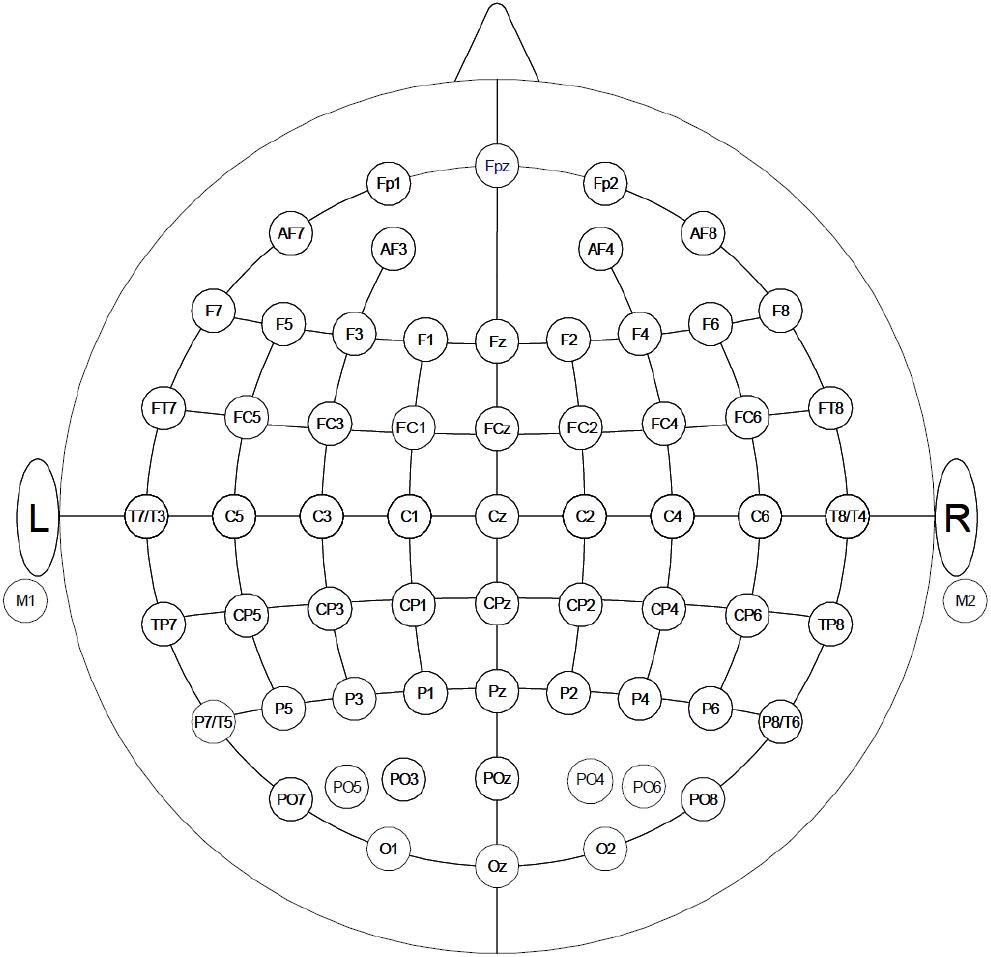
The electrode locations of EEG.

In addition, signals around the eye were recorded simultaneously with the EEG signals with two electrodes placed on the top of the right eye (referenced to the left earlobe) and next to the outer canthus of the eye (referenced to the right earlobe) to detect blinking and eye movement. The ground electrode was placed on the left wrist. All channels were amplified using a Refa 72-channel amplifier (TMSi) against the average of all connected inputs. The signals were sampled at a sampling rate of 2,048 Hz and recorded using Polybench (TMSi, EJ Oldenzaal, The Netherlands). For synchronization of auditory stimuli with biological signals and the behavioral experiment, audio and button signals were recorded by the Refa amplifier using a Dual Channel Isolation Amplifier (TMSi).

### 2.6 Data Analysis

#### 2.6.1 Labeling

The recorded EEGs were divided into two groups according to the familiarity ratings of the participants with the melodies. Trials in which participants’ scores were “Extremely familiar” and “Moderately familiar” were labeled as “familiar,” and trials in which participants’ scores were “Slightly familiar” and “not at all familiar” were labeled as “unfamiliar.” Trials with the response “Somewhat familiar” were classified as “Disagreed.”

#### 2.6.2 Preprocessing

First, the recorded EEG signal was re-referenced to the average potential of the M1 and M2 electrodes. Second, we applied a finite impulse response notch filter (50 Hz) and a bandpass filter (1–60 Hz). Third, to remove any artifacts caused by either eye movement or blinking, we applied independent component analysis (Jung et al., 2000) to the EEG and excluded independent components that were highly correlated with two-ch signals around the eye. The correlation was measured using a Pearson coefficient with a threshold of 0.9. One participant was excluded from the analysis due to unstable EEG amplitudes.

### 2.7 Event-related potential

We divided the preprocessed EEG into two groups (familiar and unfamiliar) and averaged each group. We averaged the EEGs at six regions (left frontal: F1, F3, F5; right frontal: F2, F4, F6; left central: C1, C3, C5; right central: C2, C4, C6; left parietal: P1, P3, P5; right parietal: P2, P4, P6). To compare the ERPs appearing in familiar and unfamiliar conditions, we performed a paired-sample t-test on the mean of the potentials in the three ranges (−0.5 to 0 s, 0.25 to 0.35 s, and 0.35 to 0.45 s). We set these ranges because we expected that Bereitschaftspotential (BP), P300, and N400 would appear in each interval (Wang et al., 2018; Bianco et al., 2017; Quinzi et al., 2019). We set the significance level of the t-test at *p* < 0.05.

In addition, we performed a linear regression analysis on the potentials from −0.5 to 0 s to confirm positivity of the slopes. We set the significance level in the linear regression analysis at p < 0.05.

### 2.8 Time frequency analysis

For the 4 s epoch between 2 s before and 2 s after the melody stopped, we computed the spectrogram relative to the baseline as follows:

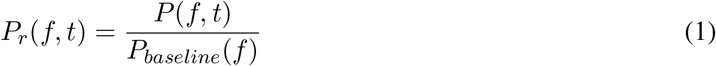

where *t* is the time defined as *t* = 0 at the start of silence, *P*(*f*, *t*) is the spectrogram of the epoch, and *P_baseline_*(*f*) is the power spectrum averaged from −2 to −1 s of the epoch. We used multitaper wavelets (Gramfort et al., 2014) for the spectrograms.

We evaluated the difference between the spectrograms of the two groups using the ratio of the spectrograms between familiar and unfamiliar groups as follows:

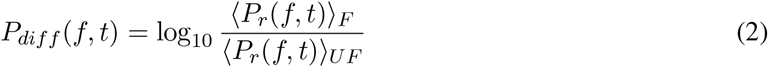

where 〈*P_r_*(*f*, *t*)〉*_F_*, 〈*P_r_*(*f*, *t*)〉*_UF_* are the epoch averages of the spectrograms within each group: familiar and unfamiliar. Then, using the ratio of the spectrograms *P_diff_* (*f*, *t*) as the statistic, we performed the cluster-based permutation test (Maris and Oostenveld, 2007), which is a nonparametric statistical test considering multiple comparisons. We set the significance level at *p* < 0.05.

### 2.9 Time-resolved multivariate pattern analysis

We conducted time-resolved multivariate pattern analysis (tMVPA) decoding of familiarity using time-series data of alpha power in melody prediction to assess temporal dynamics (Kato et al., 2022). We used the Light Gradient Boosting Machine (LightGBM) (Ke et al.) for decoding.

To perform group-level inference, we built a decoding model for each participant and combined the decoding performance of each participant. This approach is commonly used in cognitive neuroscience and facilitates individually-tailored analyses (Kragel et al., 2018; Wang et al., 2020).

We extracted the time series of the alpha power by marginalizing the spectrogram over 8–12 Hz, which was derived from the EEG downsampled to 100 Hz. This time series of 2 s was epoched into 40 segments with a window width of 200 ms (20 samples) and a shift width of 50 ms (five samples). Since the number of electrodes is 64, each epoch has 64 × 20 = 1, 280 samples.

For each epoch, random downsampling was applied to balance the familiar and unfamiliar-class labels. The balanced dataset was shuffled to divide into training and test sets at a ratio of 8 : 2. Subsequently, we normalized all sample values in the training data into the range between 0 and 1 based on the maximum and minimum values. Then, we tuned the LightGBM parameters using LightGBMTuner (Ozaki, 2020) on the training data. This process was performed 60 times independently to evaluate a one-sided Wilcoxon signed rank that indicated whether the grand mean was significantly above the chance level (50%). We set the significance level at *p* < 0.05.

### 2.10 Phase transfer entropy

We computed the directed PTE (dPTE) (Morillon and Baillet, 2017; Lobier et al., 2014; Gelding et al., 2019) to observe the connectivity between the different brain regions. First, we divided the EEG into four epochs (−1.0 to −0.5 s, −0.5 to 0 s, 0 to 0.5 s, and 0.5 to 1.0 s) where 0 s is the moment of silence. Then, dPTE was computed from the four epochs for each of the three frequency bands, theta (4–8 Hz), alpha (8–12 Hz), and beta (15–25 Hz), as divided by the band-pass filter. The dPTE was computed using the MATLAB open code PhaseTE MF.m, using the bin size (h) of Scott (2015):

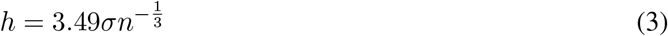

(3) where *σ* is the standard deviation of the phase data computed from the Hilbert transform for the time series, and *n* is the length of time series (number of samples). We chose four regions (left sensory-motor cortex: C3, right sensory-motor cortex: C4, left auditory cortex: T3, right auditory cortex: T4) to calculate dPTE (Gelding et al., 2019). Then, we normalized the results between −0.5 and 0.5 with the sign indicating the direction of the connectivity, as Morillon and Baillet (2017) and Gelding et al. (2019) performed. A paired-sample t-test was applied to the dPTE to confirm the difference between familiar and unfamiliar conditions. We set the significance level at *p* < 0.05.

### 2.11 Steady-state evoked potentials

To compare the frequency response of the EEG during the melody and silent intervals, we computed the averaged spectral densities of the EEG signals in the melody interval (−2 to 0 s) and the silent interval (0 to 2 s) for the two groups (familiar and unfamiliar). We also obtained the grand average, the spectral densities averaged across all electrodes and participants. We then performed two-way repeated-measures analysis of variance (ANOVA) tests for the spectral densities. Familiarity (familiar and unfamiliar) and interval (melody and silence) were independent variables, and each participant’s peak spectral density value was the dependent variable. We set the significance level at p < 0.05.

## 3 Results

### 3.1 Behavioral Analysis

We divided the recorded EEG into two groups (familiar and unfamiliar) based on the participants’ evaluation of their familiarity with the melody. The results of the familiarity evaluation are shown in Table 1.

**Table 1:** Results of familiarity assessment with the melody by participants

| Title | Familiar | Unfamiliar | Disagreed |
| --- | --- | --- | --- |
| Symphony “Ode to Joy” | 52 | 3 | 2 |
| Symphony No.9 “New World” | 50 | 2 | 5 |
| Pomp and Circumstance | 52 | 5 | 0 |
| Messiah “Hallelujah” | 29 | 22 | 6 |
| Planets “Mercury” | 38 | 15 | 4 |
| Wedding March | 55 | 1 | 1 |
| Eine Kline Nachtmusik | 50 | 5 | 2 |
| Turkish March | 56 | 0 | 1 |
| Csikos Post | 55 | 1 | 1 |
| Orpheus in Hell | 50 | 3 | 4 |
| Swan Lake “Scene” | 38 | 15 | 4 |
| The Nutcracker “March” | 52 | 3 | 2 |
| Amazing Grace | 42 | 11 | 4 |
| It’s a Small World | 55 | 1 | 1 |
| Jingle Bells | 53 | 2 | 2 |
| Twinkle Twinkle Little Star | 53 | 2 | 2 |
| We Wish You a Merry Christmas | 54 | 3 | 0 |
| When You Wish upon a Star | 42 | 15 | 0 |
| Yankee Doodle | 28 | 28 | 1 |
| B-I-N-G-O | 39 | 7 | 11 |
| The Gymnopédies | 13 | 44 | 0 |
| Piano Sonate Op.82 | 1 | 56 | 0 |
| Piano Sonate Op.14–1 | 0 | 54 | 3 |
| Sonatine Op.151-2 | 4 | 51 | 2 |
| Waltz | 0 | 56 | 1 |
| Serenade for Strings Op. 22-5 | 2 | 53 | 2 |
| Dolly Suite, Kitty-valse | 0 | 57 | 0 |
| Lyric Pieces | 0 | 53 | 4 |
| Keyboard Sonata No.12 | 1 | 56 | 0 |
| Keyboard Sonata No. 28 | 0 | 57 | 0 |
| Keyboard Sonata No. 33 | 1 | 55 | 1 |
| Sonatine Op.55-1 | 5 | 50 | 2 |
| Humoresque | 0 | 56 | 1 |
| Songs Without Words Op.19- | 2 | 53 | 2 |
| Piano Sonate KV309 | 3 | 52 | 2 |
| 10 Pieces Op.12–2 Gavotte | 0 | 56 | 1 |
| 10 Pieces Op.12–3 Rigaudon | 2 | 54 | 1 |
| Piano Sonata No.4 Scherzo | 0 | 57 | 0 |
| Piano Sonata No.6 mov.3 | 0 | 57 | 0 |
| Piano Sonata Op. 1 | 2 | 55 | 0 |

### 3.2 Event-related potential

The EEGs averaged for each of the six regions for all participants in familiar and unfamiliar groups are shown in Figure 4. As a result of t-tests in the three intervals (BP, P300, and N400) for familiar and unfamiliar groups, there were significant differences in P300 (0.25 to 0.35 s) in the right parietal and N400 (0.35 to 0.45 s) in the left frontal (right parietal: *p* < .01; *t*(18) = 4.06; *d* = 1.26, left frontal: *p* = .027; *t*(18) = −2.41; *d* = 0.61). On the other hand, there was no significant difference in BP (−0.5 to 0 s), *p* > 0.05.

**Figure 4:**
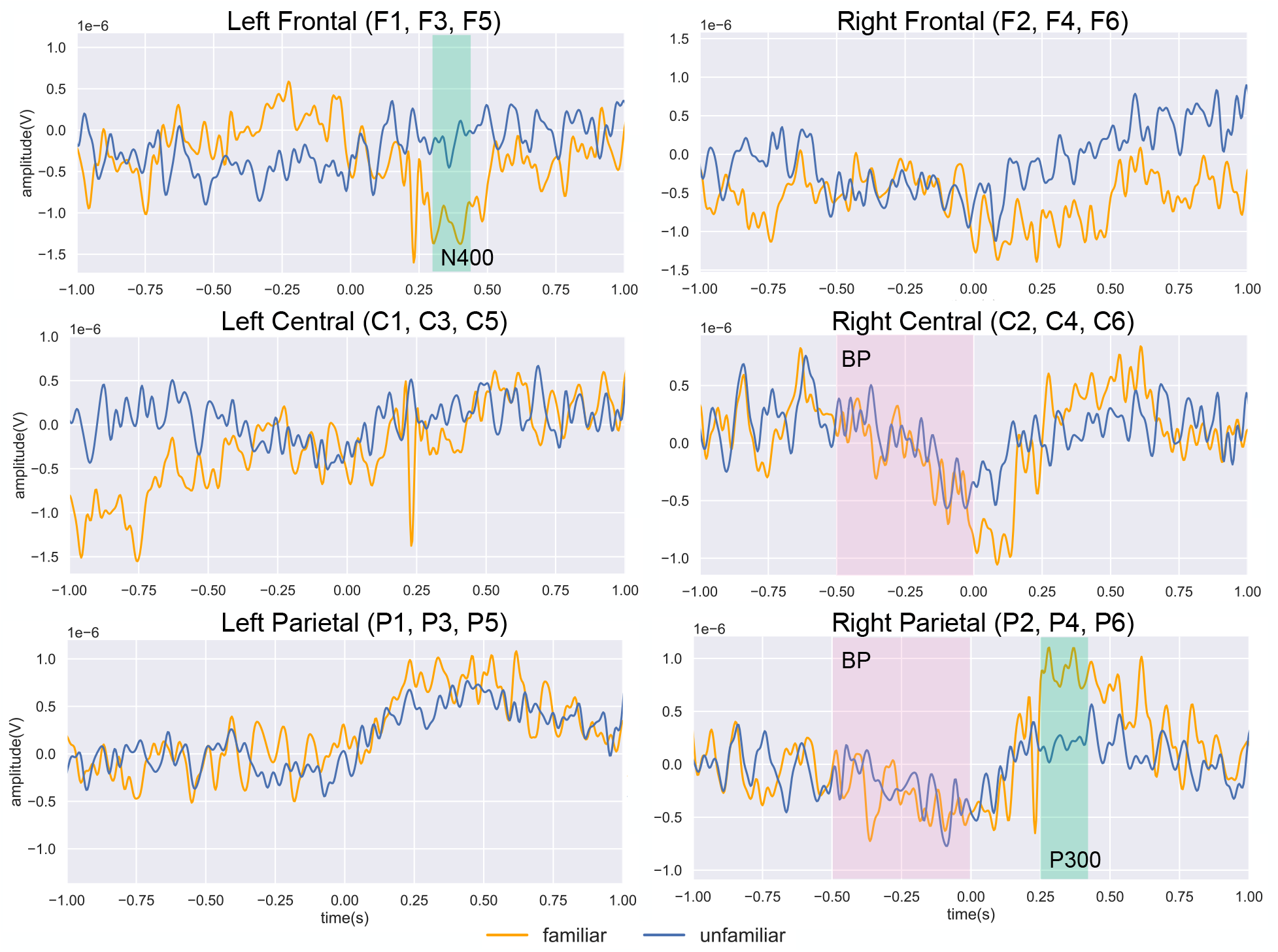
Results of averaging the EEG in the familiar (orange) and unfamiliar (blue) groups

We performed linear regression analysis on the potentials between −0.5 and 0 s (BP). As shown in Figure 4, and confirmed the significance of the negative slope in the right central area for both familiar (gradient= −1.584e −6, *t* = −14.593, *p* < .001) and unfamiliar (gradient= −1.573e −6, *t* = −16.902, *p* < .001) conditions. Also, we confirmed a significant negative slope in the right parietal area for both familiar and unfamiliar ones.

### 3.3 Time frequency analysis

The ratio of the spectrograms between familiar and unfamiliar groups is presented in Figure 5 A. As shown in Figure 5 A, alpha power was more suppressed just before silence (−0.5 s) in the left frontal (F3) and left central (C5) regions when familiar melodies were presented than unfamiliar ones.

**Figure 5:**
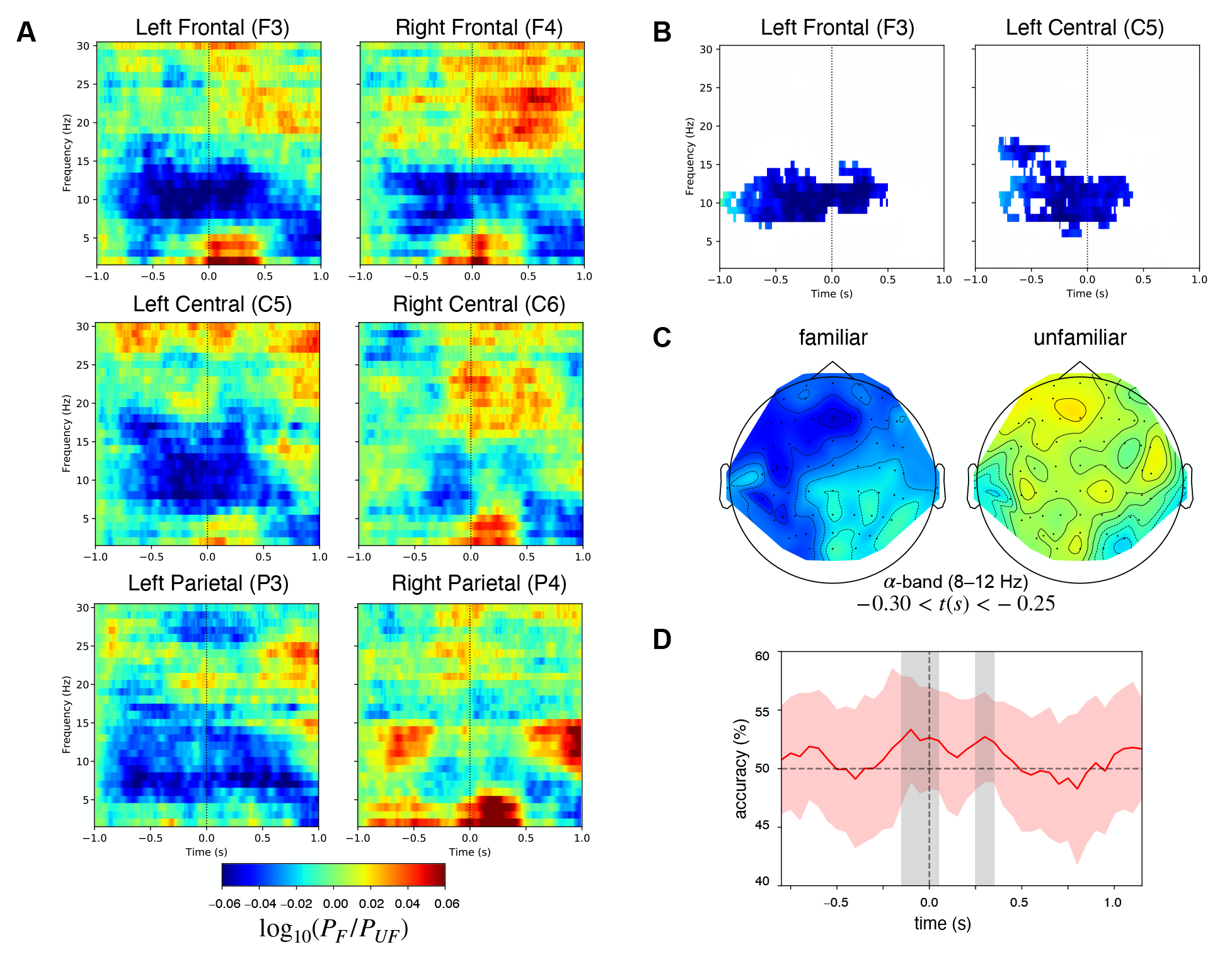
(A) Spectrograms of the familiar and unfamiliargroups. (B) Spectrograms showing only when *p* < 0.05 on the cluster-based permutation test. (C) Topographical map of alpha power (8–12 Hz). (D) The grand mean accuracy of participant-wise decoding. Gray shading indicates statistical significance (Wilcoxon signed-rank test, *p* < 0.05). Red shaded areas indicate 95 % confidence intervals across participants.

The topographical maps of alpha power averaged within each group are shown every 50 ms in Figure 5 C. In the topographical maps, alpha power was suppressed in the familiar group’s frontal to the left central region. Furthermore, in the cluster-based permutation test, we identified a significant difference between −0.5 and 0 s in the left frontal and left central regions (Figure 5 B).

The temporal variation of the accuracy rate by tMVPA is shown in Figure 5 D. To avoid the delay impact, we plotted the grand mean of the accuracy rate at the last point of each window. As shown in Figure 5 D, there were significant differences in the ranges between −0.15 and 0.05 s and between 0.25 and 0.35 s.

### 3.4 Phase transfer entropy

For the dPTE averaged within each group (familiar and unfamiliar), we plotted the connectivity between the right auditory cortex and the sensory-motor cortex (Figure 6 A and B), between the left auditory cortex and the sensory-motor cortex (Figure 6 C and D), and between hemispheres in the same region (Figure 6 E and F). As shown in Figure 6 A, in the theta band before silence (−1.0 to −0.5 s), we observed information flow from the right auditory cortex (rAUD) to the left sensory-motor cortex (lSM) when the familiar melody was presented (*p* = .040; *t*(18) = 2.210; *d* = 0.507). As shown in Figure 6 B, in the beta band before silence (−1.0 to −0.5 s), we also observed stronger information flow from the right sensory-motor cortex (rSM) to the rAUD for familiar melodies than for unfamiliar ones (*p* = .038; *t*(18) = −2.240; *d* = 0.514). Furthermore, in the theta band after silence (0 to 0.5 s), we observed information flow from the rSM to the rAUD during presentation of the familiar melody, and conversely, from the rAUD to the rSM during the unfamiliar one (*p* = .038; *t*(18) = −2.236; *d* = 0.513).

**Figure 6:**
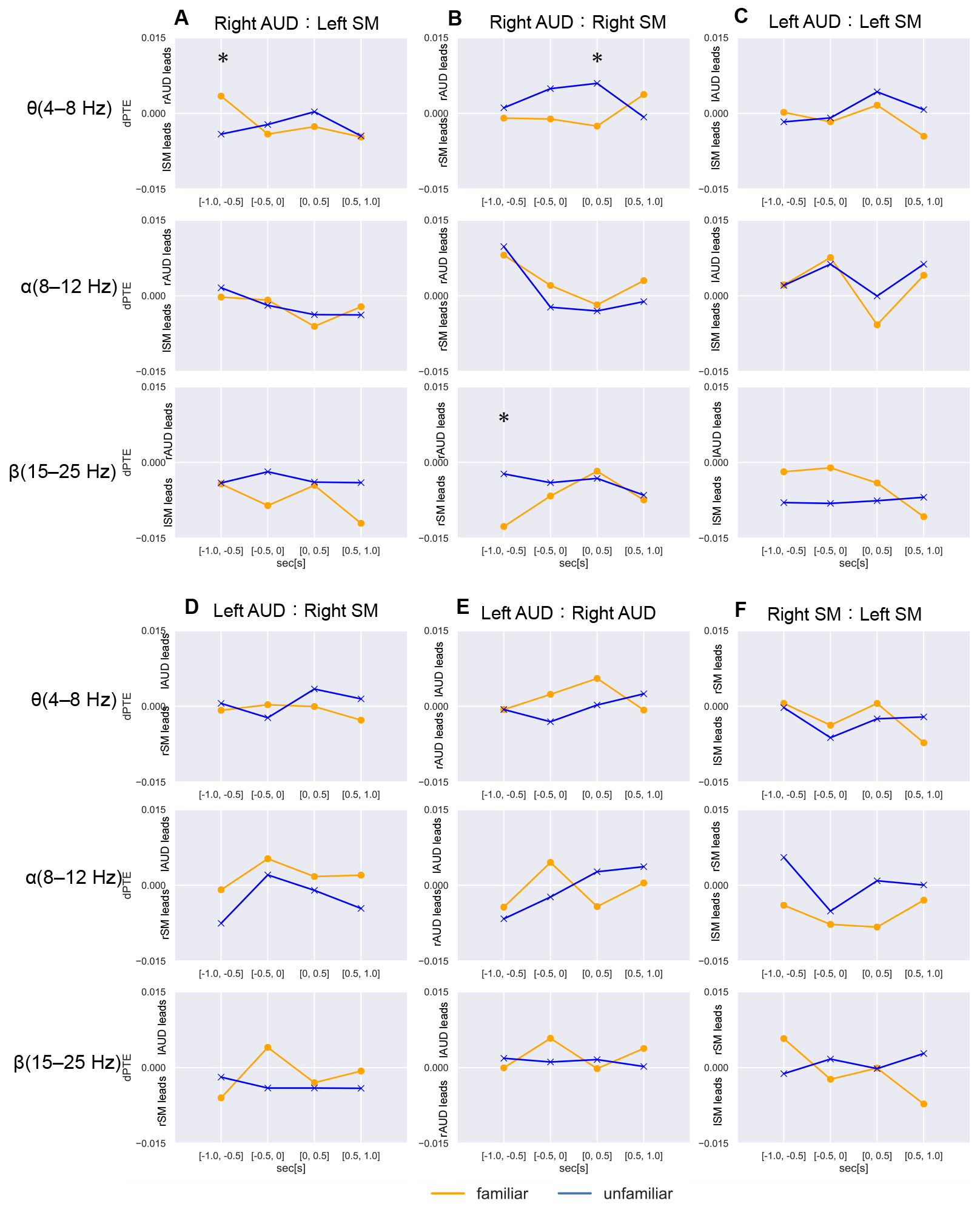
dPTE mean values for the three different frequency bands of interest, *θ*, *α*, and *β* during familiar melody (orange) and unfamiliar melody (blue) for all connections (via column): (A) rAUD to lSM; (B) rAUD to rSM; (C) lAUD to lSM; (D) lAUD to rSM; (E) lAUD to rAUD; (F) rSM to lSM. Values were computed over four equal epochs of the 2 s presentation. (A)–(D) positive values indicate the sensory-motor region activity leading the auditory region, and negative values indicate the auditory region activity leading the sensory-motor region. (E), (F) positive values indicate the left source activity leading the right source, and negative values indicate the right source activity leading the left source. * : *p* < 0.05. Abbreviations: rAUD, right auditory cortex; lSM, left sensory-motor cortex; rSM, right sensory-motor cortex; lAUD, left auditory cortex

### 3.5 Steady-state evoked potentials

We computed the spectral density of the EEG during the melody and silent intervals for familiar and unfamiliar. The spectral density of the EEG averaged across trials, all electrodes, and all participants are presented in Figure 7. As shown in Figure 7, there were peaks at multiples of the melody tempo (2.5 Hz and 5.0 Hz) in the melody interval for both familiar and unfamiliar music. There were no peaks in the silent interval for both familiar and unfamiliar. Furthermore, as a result of a two-way repeated-measures ANOVA test for the spectral density at 2.5 Hz and 5.0 Hz, we found a significant difference between the melody and silent intervals (*F* = 6.697, *p* = .012, 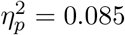).

**Figure 7:**
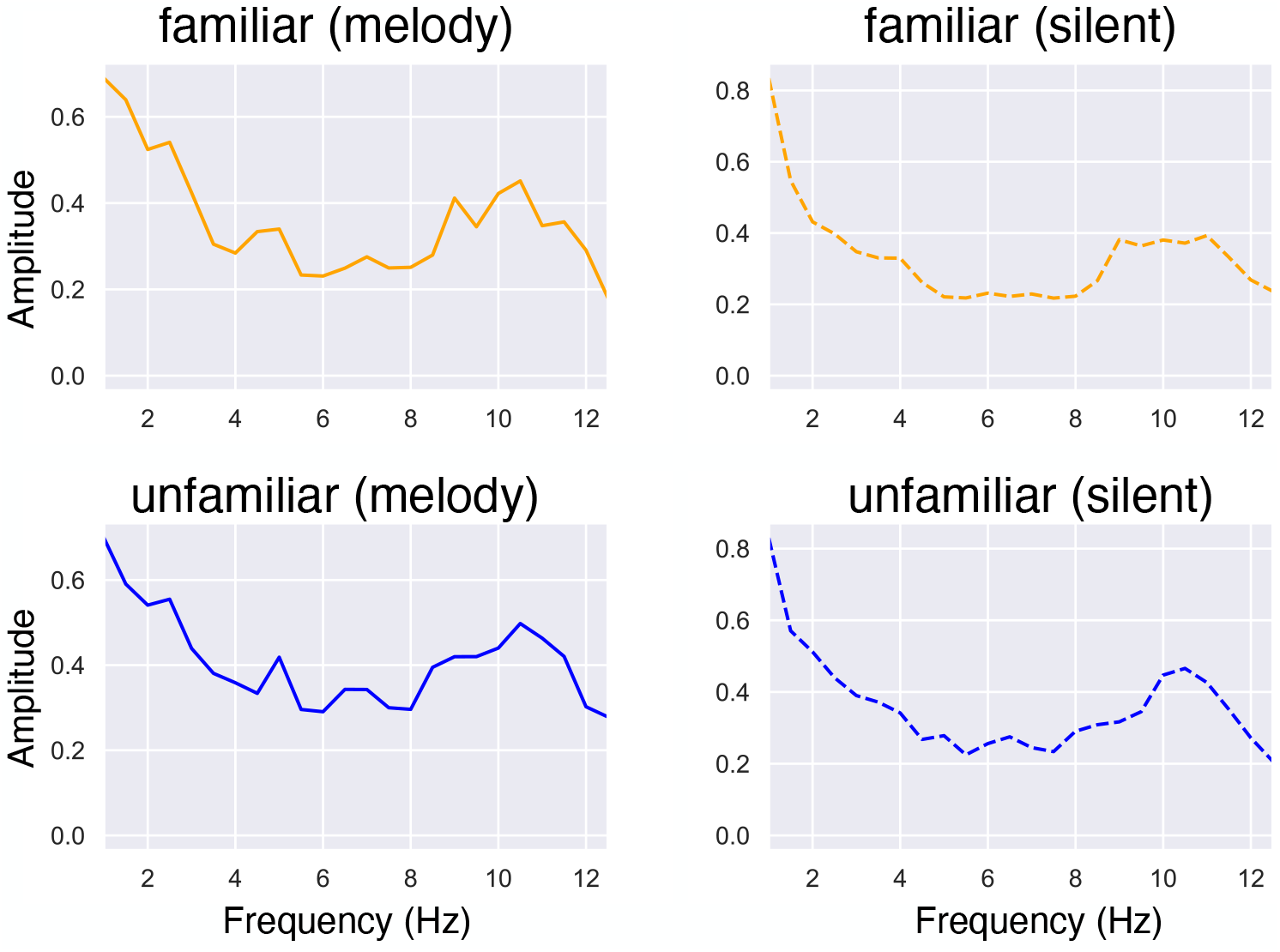
Spectral density averaged across participants. The solid orange line represents the melody interval during the familiar melody, and the solid blue line represents the melody interval during the unfamiliar melody. The dashed orange line represents the silent interval during the familiar melody, and the dashed blue line represents the silent interval during the unfamiliar melody.

## 4 Discussion

The main findings are summarized as follows: First, we found no significant difference in the EEG amplitude before silence for familiar and unfamiliar conditions. On the other hand, in the two intervals after the onset of silence (P300 and N400), there were significant differences in the right parietal and left frontal regions. Second, the regression analysis of the EEG revealed a negative slope in the central and right parietal regions before the silence. We found a significant alpha-band power suppression in the time-frequency analysis just before silence. Third, we observed stronger information flow from the right sensory-motor cortex (rSM) to the right auditory cortex in the beta band for familiar melodies than for unfamiliar ones using dPTE.

### 4.1 Brain activity associated with melody prediction

We discuss the brain activity associated with melody prediction. The significant differences in the potential from 0.25 to 0.35 s in the right parietal region and from 0.35 to 0.45 s in the left frontal region between the familiar and unfamiliar conditions (Figure 4) may be due to the appearance of P300 and N400 under familiar conditions. For the ERPs to auditory stimuli, Polich (2007) reported that P300 reflects attention after perception, and Miranda and Ullman (2007) stated that N400 reflects violation from the context. If we accept these findings, we suggest that the appearance of P300 and N400 under the familiar condition may be due to participants’ perception of silence as a violation of the melody.

In both familiar and unfamiliar conditions, we found a significant negative slope in the right central and right parietal potentials from −0.5 to 0 s. As a similar phenomenon, negative slope fluctuations were observed in the central and parietal regions before 0.5 s of stimulus presentation when external triggers controlled the auditory stimuli (Berchicci et al., 2016; Bianco et al., 2017; Quinzi et al., 2019). This is the potential for exercise preparation (BP). The BPs in the present study appeared because the participants prepared for the silence. However, it may be strange for the appearance of BP just before the silenct interval because we did not announce the silent interval in our experimental design. Regarding this issue, we provided the silence about 8 s after the start of the melody so that the participants could predict timing of the silence after a few trials. For example, in the study by Quinzi et al. (2019), stimuli were presented at intervals of 1–2 s, so participants could predict the stimuli’s timing.

### 4.2 Power fluctuations associated with melody prediction

We describe alpha power suppression just before silence based on the time-frequency analysis. Alpha power was more suppressed just before silence (−0.5 s) in the left frontal and left central regions when familiar melodies were presented than for unfamiliar ones (Figure 5 B).

Alpha power is suppressed in brain activity for language prediction when the prediction of the next word is more predictable (Wang et al., 2018). Also, alpha power suppression is related to attention and expectation of the stimulus (Klimesch et al., 1998; van Ede et al., 2012). Suppression due to expectation is often observed in the parietal regions (Schroeder and Lakatos, 2009; Uhlhaas et al., 2010).

In the present study, we observed that alpha power suppression was stronger in the frontal and central regions just before the silence. Alpha power suppression in the left frontal and central regions reflects motor preparation and execution (Alegre et al., 2004; Neuper et al., 2006; Cheyne, 2013; Piai et al., 2014). The alpha band activity in this study may also be related to motor preparation for predicting the melody during the silent interval. Furthermore, we suggested that the negative slope of the EEG just before the silent interval might be due to BP, and the alpha power suppression supports this suggestion.

In addition, there was a tendency for beta power to increase after silence in the frontal region (Figure 5 A). For this phenomenon, it is likely an increase in frontal beta-power for language prediction (Wang et al., 2018; Terporten et al., 2019). Alternatively, the beta power increase in the prefrontal cortex may be related to working memory (Schmidt et al., 2019). Thus, it may be an activity related to memory for music. More research is required to enable greater understanding of this phenomenon.

### 4.3 Relationship between auditory and sensory-motor cortex in melody prediction

This section describes the relationship between the auditory and sensory-motor cortex in melody prediction. We observed that information flows from the rSM to the rAUD was stronger in the beta band before the silence for familiar melodies than for unfamiliar ones (Figure 6 B).

Beta-band modulation is related to the prediction of timing in both perceptions (Fujioka et al., 2012) and imagery (Fujioka et al., 2015). Furthermore, Gelding et al. (2019) observed stronger beta-band modulation during silence when sounds were more predictable than when they were less predictable. For this phenomenon, beta-band modulation reflects predictability of the event (Chang et al., 2016, 2018; Gelding et al., 2019). In addition, the right hemisphere is related to the imagery of music (Lacey and Lawson, 2013; Zatorre and Halpern, 2005). Considering these findings, we suggest that the information flows from the rSM to the rAUD in the beta band before the silence reflects the timing of music imagery and prediction.

Furthermore, we observed information flow from the rSM to the rAUD in the theta band after the onset of silence when a familiar melody was presented. On the other hand, information flows in the opposite direction when an unfamiliar melody is presented (Figure 6 B). Additionally, these directions were reversed within 0.5 s (from 0.5 to 1 s) after the silence. This finding is relevant to the suggestions by Moore (2010) and Ashley and Timmers (2017), who posited two different types of imagery: an obligatory form that is invoked to assist perception (constructive imagery); and a form of sensory imagery that can be invoked voluntarily in the absence of sensory input. Gelding et al. (2019) also suggest that a reverse of information flow appears in the perception and imagery of music. Therefore, in the present study, the reverse of information flow between easily predictable (familiar) and less predictable (unfamiliar) melodies may be related to these two types of imagery.

### 4.4 Frequency response of EEG in the silent interval

This section describes the frequency response of the EEG in the silent interval. We observed peaks at multiples of the melody tempo (2.5 and 5.0 Hz) in the melody section for both the familiar and unfamiliar conditions (Figure 7). On the other hand, there were no peaks in the silent interval under either condition.

It has been reported that steady-state evoked potentials (SSEPs) appear when individuals listen to music (Nozaradan et al., 2012; Meltzer et al., 2015; Kumagai et al., 2017). In addition, Okawa et al. (2017) demonstrated that SSEPs of the same frequency as the rhythm appeared when participants imagined a periodic rhythm. Based on these results, we expected that SSEPs of tempo would appear because participants could imagine music during silence for familiar melodies. However, SSEPs appeared in the melody section but not in the silent section. This may be because the participants could not capture the rhythmic timing well during the silent intervals. Besides, we mainly used classical music as the presenting stimulus. Music with a strong beat, such as hip-hop, might induce SSEPs even during the silent interval because the participants might be able to easily maintain the rhythm.

## 5 Conclusion

We hypothesized that there are similarities between the physiological mechanisms of prediction in music and language. We measured and analyzed the EEG of participants while they listened to a melody with a silent section. For this hypothesis, we obtained similar results: alpha power suppression was observed in the left frontal and left central regions for familiar music. Besides, we identified new physiological mechanisms for music prediction; the appearance of BP in the right central and right parietal regions and the information flow from the rSM to the rAUD in the beta band. However, one of the limitations of this study is that EEG is less accurate for measuring high-frequency components, such as the gamma band (26–70 Hz), than MEG, which was employed to investigate the neural activity in language prediction with the coupling analysis between the alpha and gamma bands showing negative correlation (Wang et al., 2018). Further studies investigating the relationship between language and music prediction could use measurement methods with better frequency resolution, such as MEG and electrocorticography.

## Credit author statement

**Shuma ITO:** Conceptualization, Methodology, Software, Data Curation, Formal analysis, Validation, Investigation, Writing - Original Draft.

**Kazuki MATSUNAGA:** Software, Formal analysis, Validation.

**Ingon CHANPORNPAKDI:** Writing - Review & Editing, Investigation.

**Toshihisa TANAKA:** Conceptualization, Methodology, Validation, Writing-Reviewing and Editing, Project administration, Supervision, Resources, Funding acquisition.

## Acknowledgements

We would like to thank Editage (www.editage.com) for English language editing.

